# Exploring Log-Likelihood Scores for Ranking Antibody Sequence Designs

**DOI:** 10.1101/2024.10.07.617023

**Authors:** Talip Uçar, Cedric Malherbe, Ferran Gonzalez

## Abstract

Generative models trained on antibody sequences and structures have shown great potential in advancing machine learning-assisted antibody engineering and drug discovery. Current state-of-the-art models are primarily evaluated using two categories of in silico metrics: sequence-based metrics, such as amino acid recovery (AAR), and structure-based metrics, including root-mean-square deviation (RMSD), predicted alignment error (pAE), and interface predicted template modeling (ipTM). While metrics such as pAE and ipTM have been shown to be useful filters for experimental success, there is no evidence that they are suitable for ranking, particularly for antibody sequence designs. Furthermore, no reliable sequence-based metric for ranking has been established. In this work, using real-world experimental data from fourteen diverse datasets, we extensively benchmark a range of generative models, including LLM-style, diffusion-based, and graph-based models. We show that log-likelihood scores from these generative models have promising correlation with experimentally measured binding affinities, suggesting that log-likelihood can *potentially* serve as a reliable metric for ranking antibody sequence designs. Additionally, we scale up one of the diffusion-based models by training it on a large and diverse synthetic dataset, significantly enhancing its ability to rank antibodies based on their binding affinities. We also evaluate non–log-likelihood-based metrics on ten datasets and find that, while they are less consistent for ranking, they provide complementary information. Structure-, energy-, and sequence-based scores appear to be orthogonal and may be used together to increase the likelihood of experimental success. Our implementation is available at: https://github.com/AstraZeneca/DiffAbXL

## 1 Introduction

Antibodies are crucial components of the immune system and have become indispensable tools in therapeutics and diagnostics due to their ability to specifically recognize and bind to a wide range of antigens. Engineering antibodies to improve their affinity, specificity, and stability is a rapidly advancing field, increasingly driven by machine learning and computational approaches. Generative models trained on antibody sequences and structures hold great promise in accelerating antibody design and drug discovery. However, current state-of-the-art models typically rely on in silico evaluation metrics, divided into two primary categories: sequence-based metrics, such as amino acid recovery (AAR), and structure-based metrics, such as root-mean-square deviation (RMSD) between predicted and actual structures. Recent advances in structural prediction, notably AlphaFold [Jumper et al., 2021], have significantly improved our ability to predict protein structures. These tools provide structure-based confidence metrics, such as predicted Local Distance Difference Test (pLDDT), predicted alignment error (pAE), predicted template modeling (pTM), interface predicted template modeling (ipTM), and DockQ scores. Some of these metrics, such as pAE and ipTM, have been demonstrated to be effective filters for distinguishing between high-quality and low-quality structural models, thereby enhancing the chances of experimental success [Abramson et al., 2024, Watson et al., 2023]. While these structure-based metrics are valuable for filtering and assessing model performance, they are not suitable for ranking antibody sequence designs, particularly when it comes to predicting binding affinity and functional efficacy. Moreover, existing sequence-based metrics, such as AAR, provide limited insights into functional performance, as there is no established proxy derived from sequence information alone that accurately predicts binding affinity. This poses a substantial challenge in prioritizing antibody candidates for experimental validation.

Physics-based approaches provide energy-based metrics by modeling biological systems and accounting for factors such as protein flexibility, explicit solvents, co-factors, and entropic effects. However, the correlation between these metrics and experimentally measured binding affinities is often low [Bennett et al., 2023], and there is no strong evidence that they are effective at ranking antibody designs by affinity *in a consistent manner*. These methods also face significant challenges, including high computational costs and difficulties in automation [Alford et al., 2017], which limits their utility for large-scale affinity predictions.

In this work, we address these limitations by conducting a rigorous evaluation of state-of-the-art generative models for antibody design, using fourteen diverse real-world datasets and a range of generative approaches, including Large Language Model (LLM)-style, diffusion-based, and graph-based models. We demonstrate that log-likelihood scores from these models correlate with experimentally measured binding affinities, suggesting that log-likelihood can serve as a practical metric for ranking antibody sequence designs. While the strength and consistency of this correlation vary across targets and measurement types, our findings support the utility of log-likelihood in prioritizing candidates for experimental validation. To further enhance ranking performance, we scale up a diffusion-based model by training it on a large and diverse synthetic dataset, resulting in improved correlations with binding affinity and outperforming other models. This scaled model, DiffAbXL, offers both predictive accuracy and computational efficiency, strengthening its suitability for real-world antibody design workflows. Our evaluation is structured in two parts. We first assess log-likelihood-based ranking across seven initial datasets. In the second part, we examine a broader set of non–log-likelihood-based metrics using a combined set of ten datasets—three from the initial evaluation and seven additional, challenging datasets introduced in IgDesign [Shanehsazzadeh et al., 2023a]. While these alternative metrics are generally less consistent for ranking, they appear to provide complementary signals. Structure-, energy-, and sequence-based metrics may capture distinct aspects of antibody functionality, suggesting potential for future exploration of integrated scoring strategies. By linking generative model outputs to experimental outcomes and evaluating a diverse set of metrics, our study offers a practical framework for antibody candidate prioritization and sets the stage for more robust in silico design pipelines in therapeutic antibody discovery.

## 2 Related Work

The application of deep learning to antibody and protein design has garnered significant attention in recent years, driven by advancements in natural language processing (NLP) and geometric deep learning. These generative models can be categorized into three broad approaches: LLMs, graph-based methods, and diffusion-based methods. Additionally, they can be distinguished by their input-output modalities: sequence-to-sequence, structure-to-sequence (inverse folding), sequence-structure co-design, and sequence-structure-to-sequence frameworks. Below, we review diffusion-based methods for protein and antibody design, with an extended discussion of LLMs and graph-based approaches provided in Section D of the Appendix.

### Diffusion-based approaches

Diffusion-based models have recently emerged as a powerful approach for antibody design. These models generate new sequences and/or structures by simulating a process that progressively refines noisy input into coherent output, holding promise for capturing intricate dependencies in complex biological systems, such as protein folding dynamics and molecular interactions, over multiple iterations [Abramson et al., 2024, Jing et al., 2024]. Moreover, they have proven effective in antibody design due to their ability to handle geometric and structural constraints. Luo et al. [2022] introduced a diffusion model, DiffAb, that integrates residue types, atom coordinates and orientations to generate antigen-specific CDRs, incorporating both sequence and structural information. More recently, Martinkus et al. [2024] proposed AbDiffuser, a diffusion-based model that incorporates domain-specific knowledge and physics-based constraints to generate full-atom antibody structures, including side chains. Another recent approach, AbX [Zhu et al.], is a score-based diffusion model with continuous timesteps, which jointly models the discrete sequence space and the SE(3) structure space for antibody design.

### Our contribution

The correlation between likelihood and binding affinity is previously observed in works such as [Shanehsazzadeh et al., 2023b], where the authors used their inverse folding model IgMPNN to design antibodies targeting HER2. However, the observation was solely based on computing the percentage of binders in sampled antibody library, where higher percentage of binders in a smaller library is used as indication to draw the conclusion. More broadly, zero-shot evaluations show that log-likelihood scores can rank general protein sequences across assays [Truong Jr and Bepler, 2023], but their application to antibody binding remains underexplored. Recent work also finds such rankings unreliable across antibody fitness metrics such as binding, stability, and expression [Chungyoun et al., 2024]. In this work, i) We show the direct correlation between log-likelihood and binding affinity and we do so by conducting the same experiment across fourteen datasets, proving its generaliazibility; ii) We conduct the experiments across different types of generative models, and show their applicability in ranking antibody sequences.; iii) Building on the diffusion-based approaches, we extend one of the existing models, DiffAb [Luo et al., 2022], by training it on a large and diverse synthetic dataset, as well as a small dataset of experimentally determined antibody structures. This scaling significantly enhances the model’s ability to predict and rank antibody designs based on binding affinities, addressing one of the key challenges in antibody design: *ranking*. By incorporating experimental validation, we demonstrate that log-likelihood scores from this scaled-up DiffAb model correlate well with experimentally measured binding affinities, positioning it as a robust tool for antibody sequence design and ranking. Our work contributes to the growing body of research to move beyond simple filtering and towards effective ranking of designs based on experimental success.

## 3 Method

In this work, to include a scaled version of a diffusion-based generative model, we adapted the diffusion modelling approach, DiffAb, proposed by Luo et al. [2022]. By scaling, we mean increasing the total input sequence length up to 450 residues, as well as expanding the dataset by orders of magnitude, while keeping the model architecture parameters mostly unchanged. Similar to other works in the literature, we trained it on designing CDR3 of the heavy chain (HCDR3) of the antibody as it contributes the most to the diversity and specificity of antibodies [Jin et al., 2022, Xu and Davis, 2000, Zhou et al., 2024]. We refer to this model as DiffAbXL-H3. We also trained another version for designing all six CDRs (DiffAbXL-A). Additionally, we evaluated two more approaches: applying Polyak-Ruppert averaging [Polyak, 1990, Ruppert, 1988] to four checkpoints from the training trajectory of DiffAbXL-A to create DiffAbXL-A-Polyak, and forming a naive ensemble (DiffAbXL-A-Ensemble) by averaging the log-likelihoods of these four checkpoints. The latter two methods were compared to assess the effectiveness of Polyak averaging versus a naive ensemble. A pseudocode for Polyak-Ruppert averaging is provided in Algorithm 2 in the Appendix.

### 3.1 DiffAbXL

#### Data representation

We represent the *i*^*th*^ amino acid in a given input 𝒱 by its type *s*_*i*_ ∈ {ACDEFGHIKLMNPQRSTVWY}, *C*_*α*_ coordinate ***x***_*i*_ ∈ ℝ ^3^, and orientation ***O***_*i*_ ∈ SO(3), where *i* = 1, 2, …, *N* and *N* is the total number of amino acids in 𝒱. An input 𝒱 consists of one or more masked regions ℳ, which undergo a diffusion process, and the remaining unmasked regions 𝒰, which serve as context. Here, 𝒰 is the union of the context regions of the antibody and the antigen, where the antigen is optional, such that 𝒱 = ℳ ⋃ 𝒰. If multiple regions are masked, ℳ refers to the set of masked regions (e.g., all six CDR regions on the heavy and light chains), and 𝒰 refers to the remaining unmasked regions. Each masked region ℳ_*k*_ has *m*_*k*_ amino acids at indexes *j*_*k*_ = *l*_*k*_ + 1, …, *l*_*k*_ + *m*_*k*_, where *k* indexes the masked regions. The generation task is defined as modeling the conditional distribution *P* (*ℳ*|*𝒰*), where 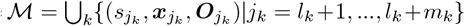 is the set of regions to be generated, conditioned on the context 𝒰 = {(*s*_*i*_, ***x***_*i*_, ***O***_*i*_)|*i* ∈ {1, …, *N*}\ ⋃_*k*_{*l*_*k*_ + 1, …, *l*_*k*_ + *m*_*k*_}}.

#### Diffusion Process

Training a diffusion probabilistic model consists of two interconnected Markov chains, referred as forward and reversed diffusion, each governing a distinct diffusion process. The forward diffusion process incrementally introduces noise into the data, ultimately approximating the prior distribution. Conversely, the generative diffusion process initiates from the prior distribution and iteratively refines it to produce the desired data distribution.

#### Forward diffusion

Starting from time *τ* = 0, noise is incrementally introduced into the data, ultimately approximating the prior distribution at time step *τ* = *T*. We use the multinomial 𝒞 (·), Gaussian (·) ∈ ℝ^3^, and isotropic Gaussian distribution ℐ 𝒢 ∈ SO(3) to add noise to the type, position, and orientation of amino acids, respectively:

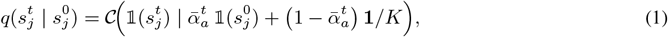

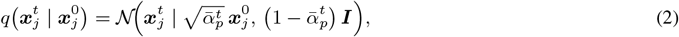

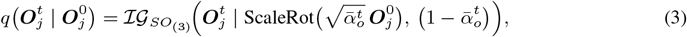

where 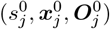 denotes the type, initial position, and orientation of the *j*^*th*^ amino acid in one of the masked regions ℳ, while 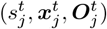 refers to their values with added noise at time step *τ* = *t*. Moreover, 𝟙 refers to one-hot encoding of amino acids, **1** is a twenty-dimensional vector filled with ones, ***I*** is the identity matrix and *K* is the total number of amino acid types (i.e., 20 in our case). In 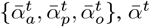 is defined as 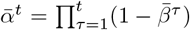, where 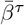 is the noise schedule for type 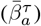, position 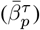, and orientation 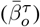 of amino acids in each masked region of ℳ at a given time *τ*.

#### Reverse diffusion

For the forward diffusion processes above, we define the corresponding reverse diffusion process as follows:

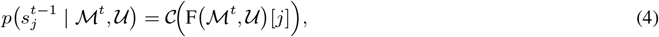

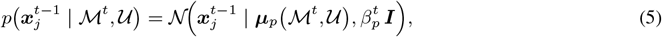

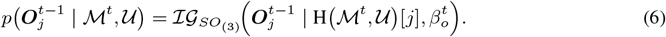

where 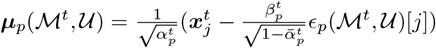, and we use F(·)[*j*], εϵ_*p*_(·)[*j*], and H(·)[*j*] to predict the type, the standard Gaussian noise _*j*_ for the position, and the denoised orientation matrix of amino acid *j* in each masked region ℳ.

#### Objective function

The training objective is defined as the sum of three losses:

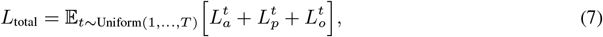

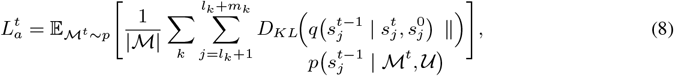

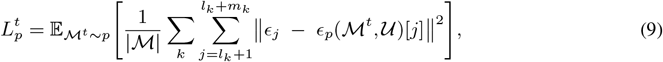

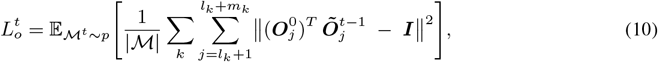

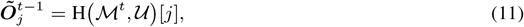

where *ϵ*_*j*_ is a standard Gaussian noise applied to the position ***x***_*j*_ of the *j*^*th*^ amino acid, and the summations over *k* account for each masked region ℳ_*k*_ that contains *m*_*k*_ amino acids indexed by *j* = *l*_*k*_ + 1, …, *l*_*k*_ + *m*_*k*_. The objective functions help the model accurately reconstruct amino acid types, positions, and orientations from noisy data. 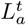 ensures correct amino acid type predictions by comparing true and predicted distributions using KL divergence. 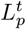 minimizes the difference between predicted and actual noise in positions, restoring spatial coordinates. 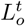 aligns the predicted and actual orientation matrices by comparing their product with the identity matrix. Together, these losses train the model to recover the masked regions consistently and accurately. Finally, for an exhaustive explanation of diffusion processes, we refer the reader to the seminal works of [Sohl-Dickstein et al., 2015, Ho et al., 2020], where the main aspects of diffusion models, including their theoretical foundations and practical applications, are covered.

### 3.2 Training

DiffAbXLs are trained on a combined dataset sourced from SAbDab [Dunbar et al., 2014] and approximately 1.5 million structures generated using ImmuneBuilder2 [Abanades et al., 2023] with paired sequences from the Observed Antibody Space (OAS) [Olsen et al., 2022a]. To ensure high-quality training data, we filtered the structures from the SAbDab dataset following the same procedure as [Luo et al., 2022], removing structures with a resolution worse than 4Å and discarding antibodies that target non-protein antigens. Next, we clustered antibodies from the combined dataset of OAS-paired sequences and SAbDab structures based on their HCDR3 sequences (or LCDR3 if HCDR3 does not exist in the sample), using a 50% sequence identity threshold for clustering. The training and test splits were determined based on cluster-based splitting. The test set included clusters containing the 19 antibody-antigen complexes from the test set used in [Luo et al., 2022] as well as 60 complexes from the RAbD dataset introduced in [Adolf-Bryfogle et al., 2018]. For validation, 20 additional clusters were selected, with the remainder used for training to maximize the training data. Both DiffAbXL-H3 and DiffAbXL-A share the same architecture and hyper-parameters, and are trained for 10 epochs using the AdamW optimizer with an initial learning rate of 1e-4, and a ReduceLROnPlateau scheduler. For further details on the model architecture, hyper-parameters and optimization, please refer to Sections E and E.1 of the Appendix.

### 3.3 Evaluation

For each sequence in a batch, we compute the log-likelihood of the masked region given the context by masking either all CDRs in antibodies and nanobodies or only the mutated CDRs. When determining the CDR region, we either use the union of several numbering schemes (AHo, IMGT, Chothia, Kabat) to account for variations in CDR definitions, or, if the designed CDRs extend beyond these regions, graft the designed CDRs into the parental sequence and use the grafted region for masking. Let *P*_*j*_(*s*_*j*_ | 𝒰) denote the posterior probability of amino acid *s*_*j*_ at position *j* conditioned on 𝒰. To ensure numerical stability, we compute the log probabilities as 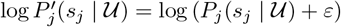, where *ε* is a small constant (e.g., 1 *×* 10^−9^). The log-likelihood for the sequence is then calculated by summing over the masked positions:

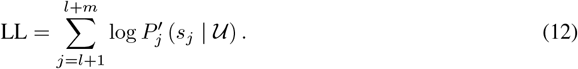

For consistency, we apply the same log-likelihood computation to BERT-style LLMs. This approach has been previously used to evaluate sequence recovery rates in works such as AntiBERTy [Ruffolo et al., 2021], AbLang [Olsen et al., 2022b], AbLang2 [Olsen et al., 2024], and IgBlend [Malherbe and Uçar, 2024]. Specifically, in this case, 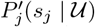 corresponds to the output of the final softmax layer at position *j* and time *τ* = 0, where the prediction is conditioned on the rest of the context 𝒰, which is provided as input to the model. Optionally, if a parent sequence is provided, with amino acids 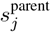, we can adjust the score by subtracting the parent’s log-likelihood:

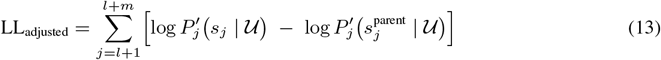

Unless otherwise specified, we use Equation 12 in our results (see Table 1). For diffusion models, we also discuss the connection of our proxy score, Equation 12, to the evidence lower bound (ELBO) and justify its usage for ranking purposes in Section F of the Appendix. After computing the log-likelihoods for all sequences, we assess their relationship with experimental labels *y*_*i*_ (e.g., binding affinities measured in the form of either −*log*(*K*_*D*_), −*log*(*IC*50), or −*log*(*qAC*50)) by computing Spearman’s rank correlation coefficient *ρ* and Kendall’s tau *τ*. The pseudocode used for computing correlations and their variance for diffusion models can be found in Section E.2 of the Appendix. Finally, for models such as DiffAbXL that use both sequence and structure at their input, we compute their scores in two modes: i) **De Novo (DN)**^2^: We mask both sequence and structure of the region and compute the log-likelihood of the sequence in the masked region at the output; ii) **Structure Guidance (SG):** We mask only the sequence, and use the structure to guide the sampling.

**Table 1:**
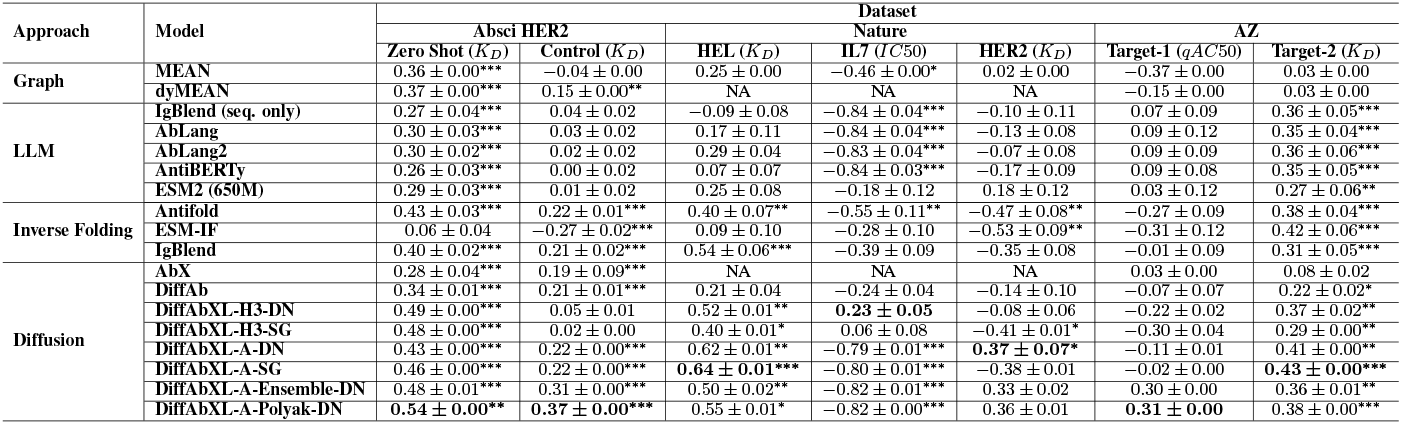
Summary of the results for Spearman correlation. Abbreviations: DN: De Novo, SG: Structure Guidance, NA: Epitope or complex structure required, but not available. *, **, *** indicate p-values under 0.05, 0.01 and 1e-4 respectively.

## 4.1 Experiments

### 4.1 Datasets

We employ fourteen datasets from four sources—Absci HER2 [Shanehsazzadeh et al., 2023b], IgDesign [Shanehsazzadeh et al., 2023a], Nature [Porebski et al., 2024], and AstraZeneca (AZ).

**The Absci HER2 datasets** [Shanehsazzadeh et al., 2023b] focus on re-designed heavy chain complementarity-determining regions (HCDRs) of the therapeutic antibody Trastuzumab, targeting HER2. The HCDRs were generated using a two-step procedure: i) CDR loop prediction using a machine learning model conditioned on the HER2 antigen backbone structure from PDB:1N8Z (Chain C), Trastuzumab’s framework sequences, and the Trastuzumab-HER2 epitope. Then, antibody sequences are sampled using an inverse folding model on predicted structures. HCDR3 lengths ranging from 9 to 17 residues were sampled based on their distribution in the OAS database while HCDR1 and HCDR2 sequences were fixed at 8 residues—common lengths for these regions. The affinity (*K*_*D*_) values of these generated sequences were measured using a Fluorescence-activated Cell Sorting (FACS)-based ACE assay. Two datasets are published: (1) the “zero-shot binders” dataset, comprising 422 HCDR3 sequences, from which we utilize those with a HCDR3 length of 13 residues (matching Trastuzumab), and (2) the SPR control dataset, which contains binders and non-binders with varying HCDR regions.

**IgDesign datasets [Shanehsazzadeh et al**., **2023a]** evaluate sequences generated for seven target antigens: FXI, IL36R, C5, TSLP, IL17A, ACVR2B, TNFRSF9. For each target, sequences were designed by mutating HCDR3 or all three CDRs of the heavy chain. The designed libraries were synthesized and experimentally evaluated for binding via SPR assays, which included appropriate positive and negative Fab controls, as well as monoclonal antibody controls.

**The Nature datasets**, published by Porebski et al. [2024], provide experimental results for three targets: HER2, HEL, and IL7. For HER2, mutations are present solely in the HCDR3 region, while for IL7, mutations occur in both LCDR1 and LCDR3 regions. In contrast, the HEL dataset consists of nanobodies with mutations across all three CDR regions. These datasets contain 25, 19, and 38 data points for HER2, IL7, and HEL respectively. We use *IC*50 measurement for IL7 and *K*_*D*_ for HER2 and HEL. Additionally, for models that require structural inputs, we predicted the structures using the parental sequences for HER2, IL7 and HEL by using ImmuneBuilder2, IgFold, and NanoBodyBuilder2 respectively [Abanades et al., 2023, Ruffolo et al., 2023] — estimated errors are shown in Table 6 of the Appendix.

**Table 2:** Comparing DiffAbXL-A-DN, Boltz-1, Rosetta, DG-Affinity and AlphaBind. Abbreviations: DN: De Novo, NA: Not Available. *, **, *** indicate p-values under 0.05, 0.01 and 1e-4 respectively.

| Model | HER2 Zero Shot | HER2 SPR Control | AZ Target-2 | IL17A | ACVR2B | FXI | TNFRSF9 | IL36R | C5 | TSLP |
| --- | --- | --- | --- | --- | --- | --- | --- | --- | --- | --- |
| DiffAbXL-A-DN | 0.43*** | 0.22*** | 0.41** | 0.62** | 0.54* | 0.18 | 0.18 | 0.14 | -0.32 | -0.02 |
| Boltz-1 ipddt | 0.26** | 0.30*** | NA | 0.38 | 0.53 | -0.05 | 0.45** | 0.23 | -0.20 | -0.22* |
| Boltz-1 plddt | 0.26** | 0.31*** | NA | 0.38 | 0.53 | -0.23 | 0.38* | 0.19 | -0.52** | -0.18* |
| Boltz-1 iptm | 0.19** | 0.12* | NA | 0.50 | 0.35 | 0.02 | 0.48** | 0.20 | 0.28 | -0.17 |
| DG-Affinity | 0.31*** | 0.49*** | NA | -0.46 | -0.11 | 0.29* | -0.27 | -0.35* | -0.17 | -0.33*** |
| AlphaBind | -0.16* | 0.03*** | -0.06 | 0.03 | -0.01 | -0.29* | 0.39* | 0.28 | -0.02 | 0.08 |
| Rosetta | 0.38*** | 0.51*** | 0.10 | 0.06 | -0.26 | 0.33* | 0.45* | 0.28 | NA | -0.04 |

**Table 3:** Comparing structure- and sequence-based metrics for binder versus non-binder classification. NA: Not Available.

| Table Type | HER2 Zero Shot | HER2 SPR Control | IL17A | ACVR2B | FXI | TNFRSF9 | IL36R | C5 | TSLP |
| --- | --- | --- | --- | --- | --- | --- | --- | --- | --- |
| **Dataset Statistics** |  |  |  |  |  |  |  |  |  |
| Dataset size | 201 | 1266 | 179 | 171 | 116 | 158 | 177 | 198 | 176 |
| KD measurements (% binders) | 201 (100%) | 420 (33%) | 13 (7%) | 21 (12%) | 47 (41%) | 35 (22%) | 40 (23%) | 34 (17%) | 127 (72%) |
| **AUC-ROC** |  |  |  |  |  |  |  |  |  |
| Boltz-1 ipddt | NA | 0.73 | 0.93 | 0.74 | 0.59 | 0.80 | 0.74 | 0.80 | 0.72 |
| Boltz-1 plddt | NA | 0.77 | 0.90 | 0.76 | 0.49 | 0.77 | 0.72 | 0.72 | 0.76 |
| Boltz-1 iptm | NA | 0.69 | 0.89 | 0.80 | 0.66 | 0.88 | 0.72 | 0.76 | 0.83 |
| DiffAbXL-A-DN | NA | 0.71 | 0.60 | 0.54 | 0.62 | 0.74 | 0.52 | 0.33 | 0.87 |

**Table 4:**
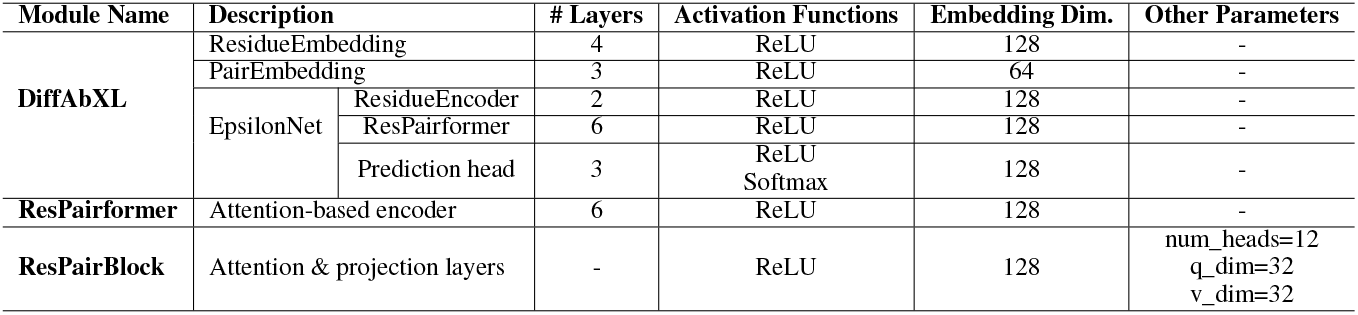
Summary of the DiffAbXL.

**Table 5:**
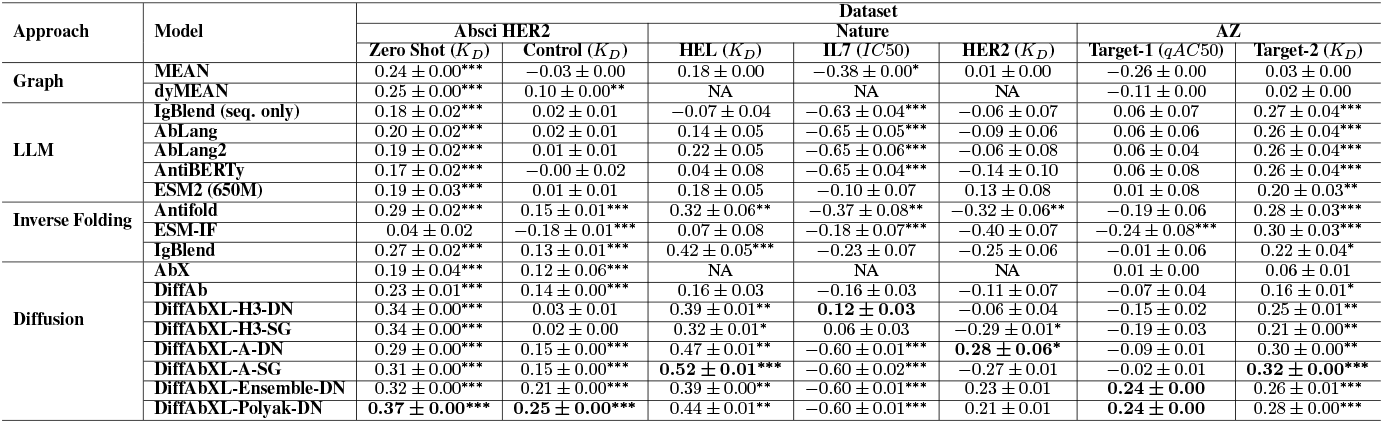
Summary of the results for Kendall correlation. Abbreviations: DN: De Novo mode, SG: Structure Guidance mode, NA: Epitope or complex structure required, but not available. *, **, *** indicate p-values under 0.05, 0.01 and 1e-4 respectively.

**Table 6:** Prediction errors for different regions of the parental nanobody used for Nature HEL and two parental antibodies used for HER2 and IL7 respectively.

| Region | Prediction Error |  |  |
| --- | --- | --- | --- |
|  | Nature HEL (Nanobody) | Nature HER2 (Antibody) | Nature IL7 (Antibody) |
| Framework H-chain | 0.87 | 0.36 | 0.37 |
| HCDR1 | 1.61 | 0.27 | 0.38 |
| HCDR2 | 1.57 | 0.36 | 0.72 |
| HCDR3 | 2.82 | 1.35 | 2.1 |
| Framework L-chain | - | 0.38 | 0.4 |
| LCDR1 | - | 0.51 | 0.92 |
| LCDR2 | - | 0.28 | 0.55 |
| LCDR3 | - | 0.30 | 0.91 |

**Table 7:** Comparison of the results with and without the antigen for Spearman correlations. Abbreviations: DN: De Novo mode, SG: Structure Guidance mode.

| Correlation | Model | Antigen | Absci HER2 |  | AZ |  |
| --- | --- | --- | --- | --- | --- | --- |
| | | | Zero Shot ($K_D$) | Control ($K_D$) | Target-1 ($qAC50$) | Target-2 ($K_D$) |
| Spearman | DiffAbXL-H3-DN | Yes | $0.49 \pm 0.00$ | $0.05 \pm 0.01$ | $-0.22 \pm 0.02$ | $0.37 \pm 0.02$ |
| | | No | $0.50 \pm 0.00$ | $-0.07 \pm 0.01$ | $-0.33 \pm 0.05$ | $0.35 \pm 0.01$ |
| | DiffAbXL-H3-SG | Yes | $0.48 \pm 0.00$ | $0.02 \pm 0.00$ | $-0.30 \pm 0.04$ | $0.29 \pm 0.00$ |
| | | No | $0.48 \pm 0.00$ | $-0.02 \pm 0.01$ | $-0.45 \pm 0.03$ | $0.29 \pm 0.01$ |
| | DiffAbXL-A-DN | Yes | $0.43 \pm 0.00$ | $0.22 \pm 0.00$ | $-0.11 \pm 0.01$ | $0.41 \pm 0.00$ |
| | | No | $0.47 \pm 0.00$ | $0.24 \pm 0.00$ | $-0.09 \pm 0.00$ | $0.31 \pm 0.02$ |
| | DiffAbXL-A-SG | Yes | $0.46 \pm 0.00$ | $0.22 \pm 0.00$ | $-0.02 \pm 0.00$ | $0.43 \pm 0.00$ |
| | | No | $0.45 \pm 0.00$ | $0.25 \pm 0.00$ | $-0.09 \pm 0.00$ | $0.41 \pm 0.00$ |
| Kendall | DiffAbXL-H3-DN | Yes | $0.34 \pm 0.00$ | $0.03 \pm 0.01$ | $-0.15 \pm 0.02$ | $0.25 \pm 0.01$ |
| | | No | $0.35 \pm 0.00$ | $-0.04 \pm 0.01$ | $-0.25 \pm 0.03$ | $0.25 \pm 0.01$ |
| | DiffAbXL-H3-SG | Yes | $0.34 \pm 0.00$ | $0.02 \pm 0.00$ | $-0.19 \pm 0.03$ | $0.21 \pm 0.00$ |
| | | No | $0.34 \pm 0.00$ | $-0.01 \pm 0.00$ | $-0.28 \pm 0.03$ | $0.21 \pm 0.01$ |
| | DiffAbXL-A-DN | Yes | $0.29 \pm 0.00$ | $0.15 \pm 0.00$ | $-0.09 \pm 0.01$ | $0.30 \pm 0.00$ |
| | | No | $0.32 \pm 0.00$ | $0.16 \pm 0.00$ | $-0.11 \pm 0.00$ | $0.22 \pm 0.01$ |
| | DiffAbXL-A-SG | Yes | $0.31 \pm 0.00$ | $0.15 \pm 0.00$ | $-0.02 \pm 0.01$ | $0.32 \pm 0.00$ |
| | | No | $0.30 \pm 0.00$ | $0.17 \pm 0.00$ | $-0.10 \pm 0.00$ | $0.30 \pm 0.00$ |

**The AZ datasets**^3^ include two distinct antibody libraries designed for two targets. The Target-1 dataset, which is based on rational design, features mutations across four regions (HCDR1-3, LCDR3), and comprises 24 data points. The Target-2 dataset consists of 85 data points and is a combination of three libraries: two rationally designed libraries (one with mutations in three heavy chain CDRs and the other with mutations in three light chain CDRs), and a third library designed using a machine learning model with mutations across all six CDRs. Finally, we use *qAC*50 for Target-1 and *K*_*D*_ for Target-2. For all AZ targets, we used their crystal structures for models that require structure.

### 4.2 Models

#### Log-likelihood-based ranking

We utilized a variety of baseline models from the literature, categorized by protein vs. antibody design, modeling approach, and input modality. For protein-based models, we included ESM2 (650M) [Lin et al., 2023], a sequence-only LLM model, and ESM-IF [Hsu et al., 2022], an inverse folding model. For antibody-specific models, we employed several sequence-only LLM models, including Ablang [Olsen et al., 2022b], Ablang2 [Olsen et al., 2024], and AntiBERTy [Ruffolo et al., 2021]. We also included IgBlend [Malherbe and Uçar, 2024], an LLM that integrates both sequence and structural information for antibody and nanobody design. In the graph-based category, we evaluated MEAN [Kong et al., 2022], and dyMEAN [Kong et al., 2023], which use sequence-structure co-design for antibodies. Among diffusion-based models, we included AbX [Zhu et al.], DiffAb [Luo et al., 2022] and our scaled version, DiffAbXL, all of which utilize sequence-structure co-design. Lastly, we assessed Antifold [Høie et al., 2023], an inverse folding model for antibodies. **Non-log-likelihood-based metrics** To complement log-likelihood-based ranking, we incorporated alternative evaluation metrics that do not rely on generative likelihoods. Specifically, we used Boltz-1 [Wohlwend et al., 2024], an open-source deep learning model that predicts biomolecular complex structures with accuracy comparable to AlphaFold3 [Abramson et al., 2024]. Boltz-1 enables ranking of designed sequences based on the confidence and plausibility of their predicted conformations. Additionally, we used Rosetta [Das and Baker, 2008] to evaluate and rank sequences using physics-based energy calculations, DG-Affinity [Yuan et al., 2023], a deep learning model trained to directly predict binding affinities using protein language model features, and AlphaBind [Agarwal et al., 2024], which also leverages protein language models and is pretrained on large-scale binding data. These approaches allow for comparative analysis across structure- and affinity-based criteria, offering orthogonal assessments to sequence likelihoods.

### 4.3 Results

#### Benchmarking LLMs, diffusion, and graph models

We evaluated a broad range of generative models, including LLM-based, diffusion-based, and graph-based models, on seven real-world datasets.

The datasets measured binding affinity in terms of *K*_*D*_, *qAC*50, or *IC*50. Our primary goal was to assess the correlation between the models’ log-likelihoods and the experimentally measured binding affinities. The results are summarized in Table 1 for Spearman and Table 5 of the Appendix for Kendall correlations respectively. In our experiments, several key observations emerged.

**First**, all generative models trained on antibody and/or protein data exhibited a degree of correlation between their log-likelihoods and binding affinity, although the strength of this correlation varied among models (see Table 1 in the main paper and Table 5 in Appendix). This consistent relationship between likelihood and affinity suggests that these models are capturing relevant aspects of antibody design, even when they were not specifically trained to design the libraries evaluated in this study. This is a crucial finding, as it demonstrates that the models generalize to unseen targets with varying success rates. However, we note that in cases where the target is entirely out of the model’s training distribution, the correlation may diminish or disappear. **Second**, the models’ log-likelihood scores retain predictive power even when synthetic structures are used as input, as demonstrated with the Nature HEL, HER2, and IL7 datasets in Table 1. **Third**, for models capable of leveraging epitope information, such as DiffAbXL, we observed only slight variations in correlation when experiments were repeated with and without the antigen as input (see Table 7 in the Appendix). This suggests that including antigen information may not substantially enhance the predictive performance of these models in certain cases. Moreover, structure-based models seem to perform better at ranking than sequence-based models, highlighting the importance of modeling structural information in antibody design. **Fourth**, we observed that a model trained specifically to redesign the HCDR3 region of the antibody (i.e., DiffAbXL-H3) is capable of evaluating sequences with mutations outside the HCDR3 region and demonstrates a strong correlation with measured binding affinity. **Fifth**, models primarily trained on one type of data (e.g., proteins or antibodies) can effectively evaluate sequences from a different data type (e.g., nanobodies in Nature HEL), showing a strong correlation with experimentally measured binding affinity. **Sixth**, we observed strong negative correlations in some experiments involving few targets, particularly when the binding affinity is measured in terms of *IC*50 and *qAC*50 (see Nature IL7 and AZ Target-1 results in Table 1). The reasons for these negative correlations are not fully understood. Although the binding and inhibition assays measuring *K*_*D*_ and *IC*50 respectively are somewhat related, they are not directly equivalent. Biological complexity and assay conditions (e.g., allosteric effects) can lead to non-linear or unexpected relationships, making it challenging to explain some of the observations. **Seventh**, it is important to note that success in established in silico metrics does not necessarily translate to better correlation with experimentally measured binding affinities. A notable example is the comparison between AbX and DiffAb, where AbX demonstrates stronger performance across several in silico metrics (see Table-1 in Zhu et al.). However, DiffAb exhibits a better correlation with the actual binding affinity measurements in our analysis (see Table-1). This discrepancy suggests that while in silico metrics may capture certain aspects of antibody properties, they do not always align with the true binding affinity, which highlights the challenges in fully replicating biological complexity through computational metrics alone. **Eighth**, naively averaging the parameters of model checkpoints using Polyak-Ruppert averaging significantly boosted the correlation for certain targets, such as the HER2 dataset, while slightly degrading the correlation for others, such as the Nature HEL dataset. Overall, DiffAbXL-A-Polyak achieved good correlation on all datasets evaluated, although two were not statistically significant. Polyak-Ruppert averaging also outperformed the naive ensemble approach. **Finally**, among the evaluated models, the scaled diffusion model, DiffAbXL, consistently outperformed others across most datasets, demonstrating the highest correlation between log-likelihood and binding affinity (see Figure 1 and Table 1). Notably, when comparing the original DiffAb model to its scaled counterpart DiffAbXL, we observed a significant improvement in performance, highlighting the impact of training the diffusion model on a much larger synthetic dataset. This scaling effect underscores the importance of data diversity and volume in enhancing model generalization and accuracy in predicting binding affinity. As models are trained on larger and more diverse datasets, the correlation between log-likelihood scores and experimental affinity measurements becomes more pronounced, suggesting that scaling is a key factor in improving predictive power for antibody design.

**Figure 1.**
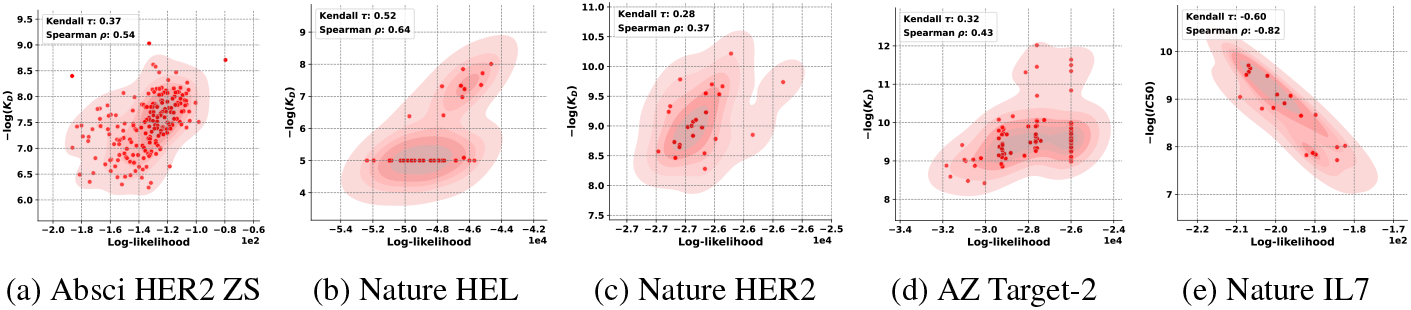
Results for DiffAbXL: **a)** DiffAbXL-A-Polyak-DN for Absci HER2 Zero-Shot (ZS), **b)** DiffAbXL-A-SG for Nature HEL, **c)** DiffAbXL-A-DN for Nature HER2, **d)** DiffAbXL-A-SG for AZ Target-2, **e)** DiffAbXL-A-Polyak-DN for Nature IL7.

#### Comparison to non-log-likelihood-based metrics

We extended our experiments to incorporate non–log-likelihood-based metrics and the datasets introduced in IgDesign [Shanehsazzadeh et al., 2023a]. Specifically, for the datasets where antigen information is available, we compared DiffAbXL’s log-likelihood scores against Boltz-1 structure-based metrics (*ipLDDT, pLDDT, ipTM*) [Wohlwend et al., 2024], Rosetta energy scores [Das and Baker, 2008], and predictions from DG-Affinity [Yuan et al., 2023], which estimates antibody binding affinities directly. The IgDesign datasets present significant challenges due to factors such as noise, limited dynamic range in *K*_*D*_, low sequence diversity, and small sample sizes. Despite these difficulties, DiffAbXL demonstrates strong and statistically significant correlations in several scenarios, as shown in Table 2. In contrast, alternative methods—including Boltz-1, DG-Affinity, Rosetta, and AlphaBind—exhibit less consistent performance, generally performing poorly except on the well-characterized HER2 target. Among these, AlphaBind performed the worst, highlighting the difficulty of predicting binding affinity even for a model specifically trained on large-scale binding data. In addition to correlation analysis, we evaluated structure- and sequence-based metrics for their effectiveness in binary classification of binders versus non-binders. Table 3 presents summary statistics of the datasets along with the AUC scores obtained using each metric. We observe that structure-based metrics may be better suited for filtering purposes, as they often yield higher AUC scores. However, this advantage varies by target and the differences are generally modest. Overall, we believe that structure-, energy-, and sequence-based metrics are largely orthogonal and can be used in combination to improve the likelihood of experimental success.

## 5 Conclusion

In this work, we demonstrated that log-likelihood scores from generative models can serve as a useful metric for ranking antibody sequence designs based on binding affinity. By benchmarking a diverse set of models—including LLM-based, diffusion-based, and graph-based approaches—across seven real-world datasets, we found consistent correlations between log-likelihood and experimentally measured affinities. The scaled diffusion model, DiffAbXL, particularly stood out by outperforming other models, highlighting the benefits of training on large and diverse datasets. Our findings underscore the potential of generative models not just in designing viable antibody candidates but also in effectively prioritizing them for experimental validation. The ability of structure-based models to outperform sequence-based ones emphasizes the importance of incorporating structural information in antibody design. We also evaluated a range of non–log-likelihood-based metrics across ten datasets, including seven challenging benchmark datasets introduced in IgDesign. While these metrics were generally less consistent for ranking, they provided complementary insights. Our results suggest that structure-, energy-, and sequence-based metrics are largely orthogonal and can be leveraged together to improve the likelihood of experimental success. Areas for further investigation include understanding the negative correlations observed in certain datasets, especially those involving *IC*_50_ and *qAC*_50_ measurements. This suggests that the relationship between log-likelihood scores and binding affinity can be complex and may vary depending on the target or measurement method. Future work should explore these nuances to refine predictive models and improve ranking accuracy. We also observe variations in correlation strength across different antibody targets, indicating that predictive performance is not uniform. We believe that these inconsistencies can be mitigated by training models on ever larger and more diverse datasets. Nonetheless, binding affinity prediction and ranking for antibodies remains an unsolved problem, and continued progress will require further methodological advances. Overall, our study provides a practical framework for leveraging generative models and complementary evaluation metrics in antibody engineering, potentially accelerating the discovery and development of next-generation therapeutic antibodies.

## Acknowledgments and Disclosure of Funding

We are grateful to Owen Vickery and Lorena Roldan Martin for running the Rosetta simulations. Special thanks to Benjamin T. Porebski for his invaluable assistance with the Nature datasets. We also appreciate the support from everyone in the MLAB program at AstraZeneca, with particular thanks to Massimo Sammito, Rebecca Croasdale-Wood, Yu Qiu and Simon Chell.

## Appendix

### A Impact Statement

This research advances the use of machine learning in antibody engineering by introducing log-likelihood as a promising metric for ranking antibody designs based on binding affinity. By demonstrating an association between model-derived scores and real-world experimental data, this work provides a pathway to accelerate therapeutic antibody discovery while minimizing costly trial-and-error experimentation. Ethical and societal impacts, such as improved healthcare outcomes and broader accessibility of life-saving treatments, mirror the established considerations in the broader field of machine learning-driven drug discovery.

### B License Information

IgDesign datasets [Shanehsazzadeh et al., 2023a] are released under MIT license. Absci Her2 datasets [Shanehsazzadeh et al., 2023b] are released under BSD License. SAbDab and OAS datasets are available under a CC-BY 4.0 license. We will release our code upon the acceptance of our paper with Apache 2.0 license.

### C Background on Antibodies

Human antibodies are classified into five isotypes: IgA, IgD, IgE, IgG, and IgM. This work focuses on IgG antibodies—Y-shaped glycoproteins produced by B-cells and nanobodies, which are single-domain antibody fragments (see Figure 2a for reference). Hereafter, “antibody” refers specifically to IgG antibodies. Antibodies have regions with distinct immune functions. The Fab (fragment antigen-binding) region, comprising variable (V) and constant (C) domains from both heavy and light chains, binds antigens. Within this region, the variable domains (VH and VL) form the antigen-binding site and determine specificity. The Fv (fragment variable) region is the smallest unit capable of antigen binding, consisting only of VH and VL without constant domains. Within variable domains are framework regions and complementarity-determining regions (CDRs). Framework regions maintain structural integrity, while CDRs—three loops on both VH and VL—directly bind antigens and are crucial for specific recognition. The Fv region, essential for antigen recognition, lacks the effector functions of the full antibody. The Fab region, including both variable and constant domains, is more stable and has higher antigen affinity. The Fv region is simpler and easier to engineer for applications such as single-chain variable fragment (scFv) antibodies. The Fc (fragment crystallizable) region at the antibody’s base regulates immune responses by interacting with proteins and cell receptors. Nanobodies are compact, single-domain antibodies derived from heavy-chain-only antibodies found in animals such as camels and llamas. Smaller than traditional Fv regions, they retain full antigen-binding capacity and offer increased stability and easier production, making them valuable in therapeutic and diagnostic applications.

**Figure 2.**
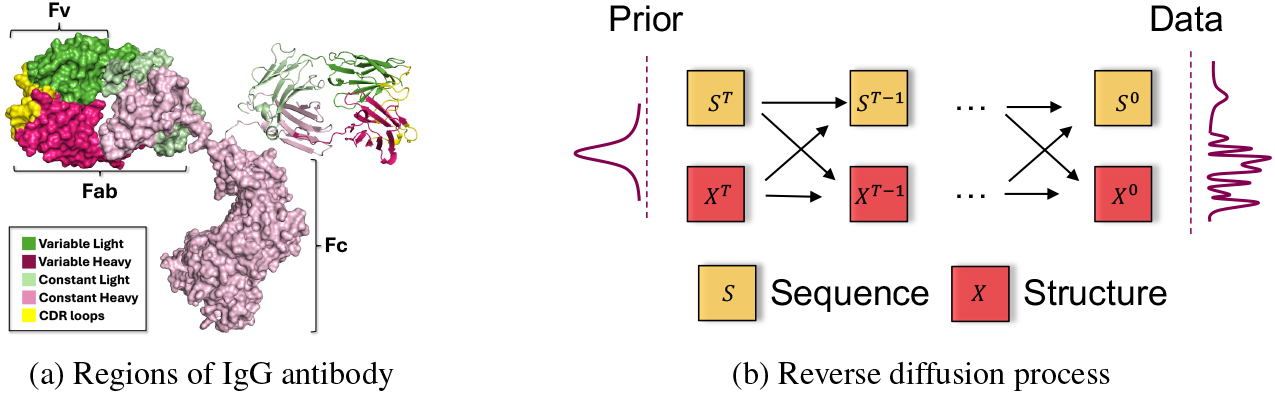
**a)** Regions of IgG antibody (PDB ID: 1igt) shown as surface (left and bottom) and cartoon (right). **b)** Iterative reverse diffusion process for DiffAbXL shown only for the position—Gaussian distribution 𝒩 (·) ∈ ℝ^3^.

**Figure 3.**
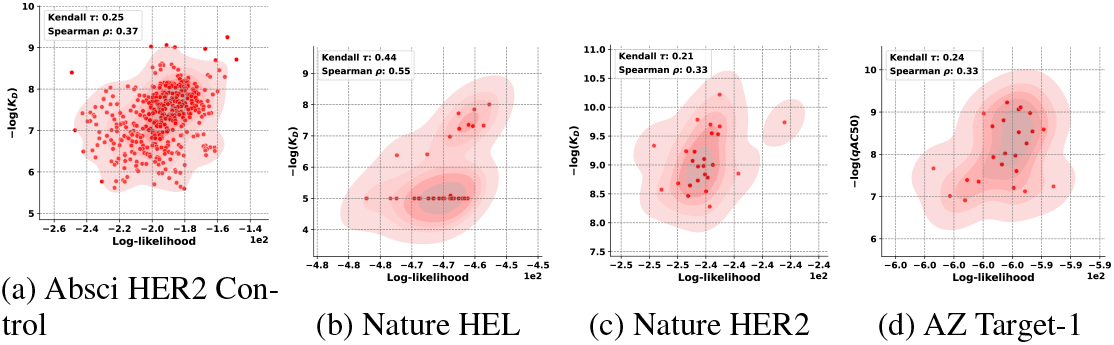
More results from DiffAbXL-A-Polyak: **a)** Absci HER2 Control, **b)** Nature HEL, **c)** Nature HER2, **d)** AZ Target-1

**Figure 4.**
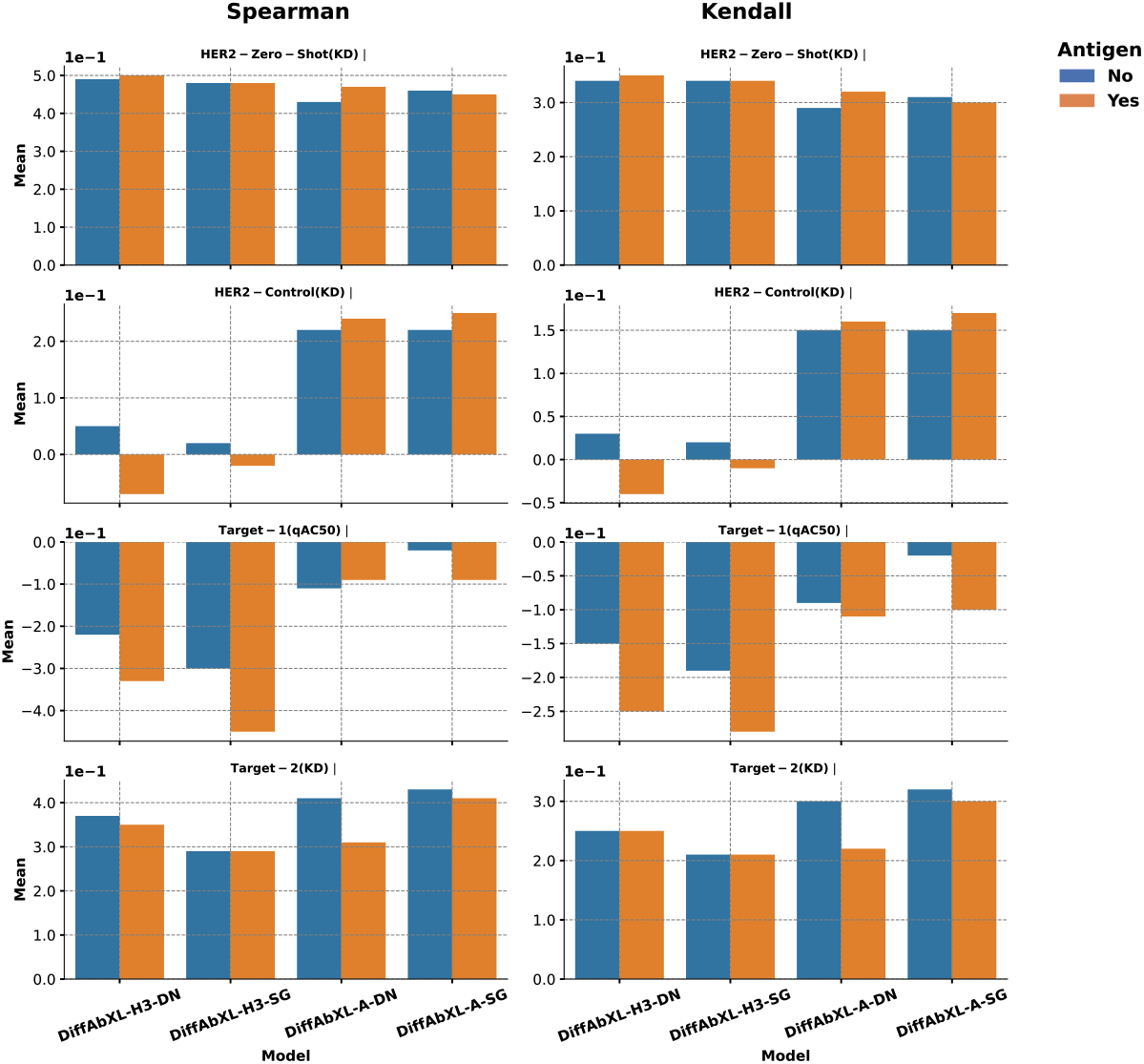
Results in Table 7 are shown as bar charts.

**Figure 5.**
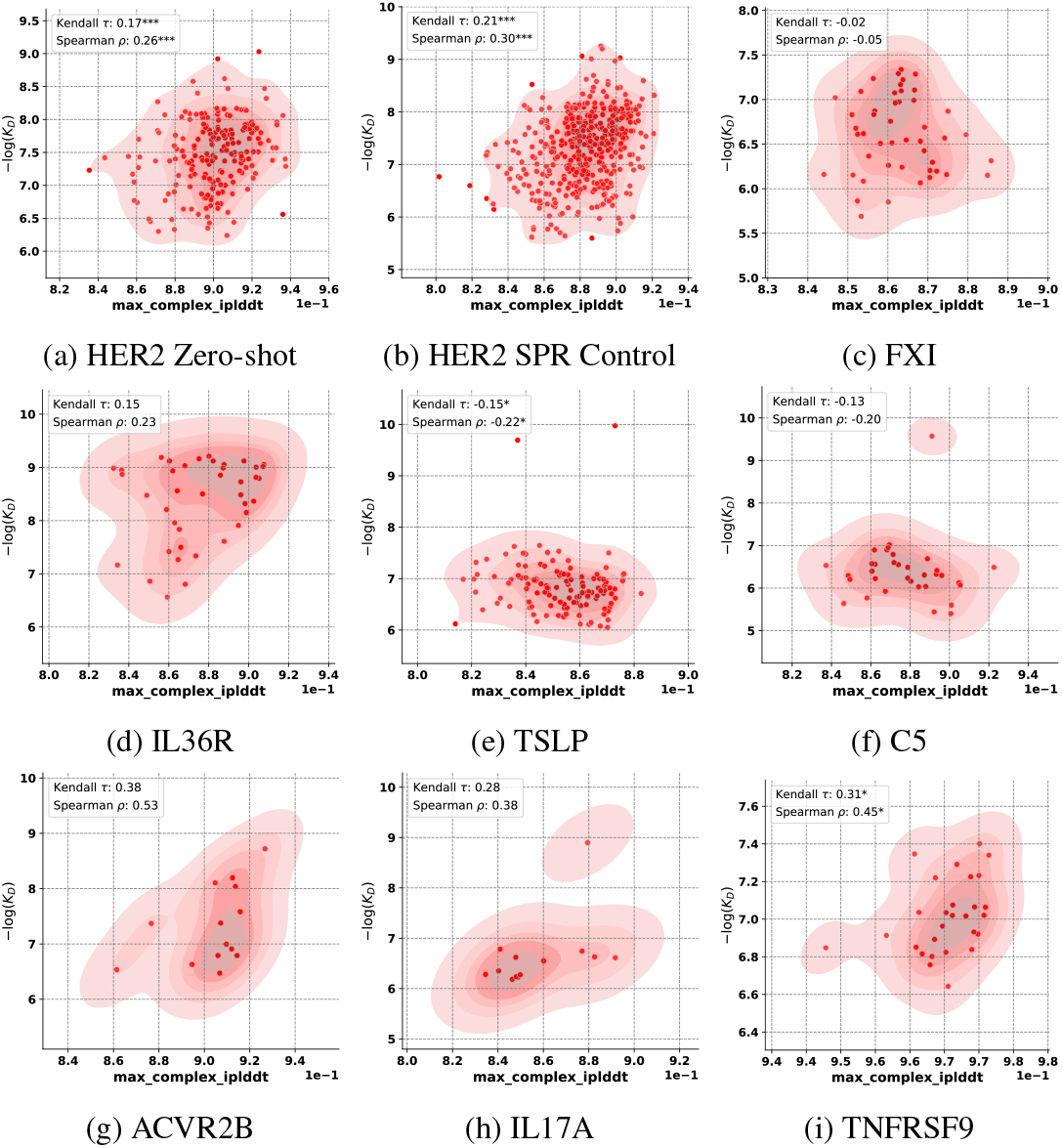
Boltz-1 Results using ipLDDT: **a)** HER2 Zero Shot, **b)** HER2 SPR Control, **c)** FXI, **d)** IL36R, **e)** TSLP, **e)** C5, **e)** ACVR2B, **e)** IL17A, **e)** TNFRSF9

**Figure 6.**
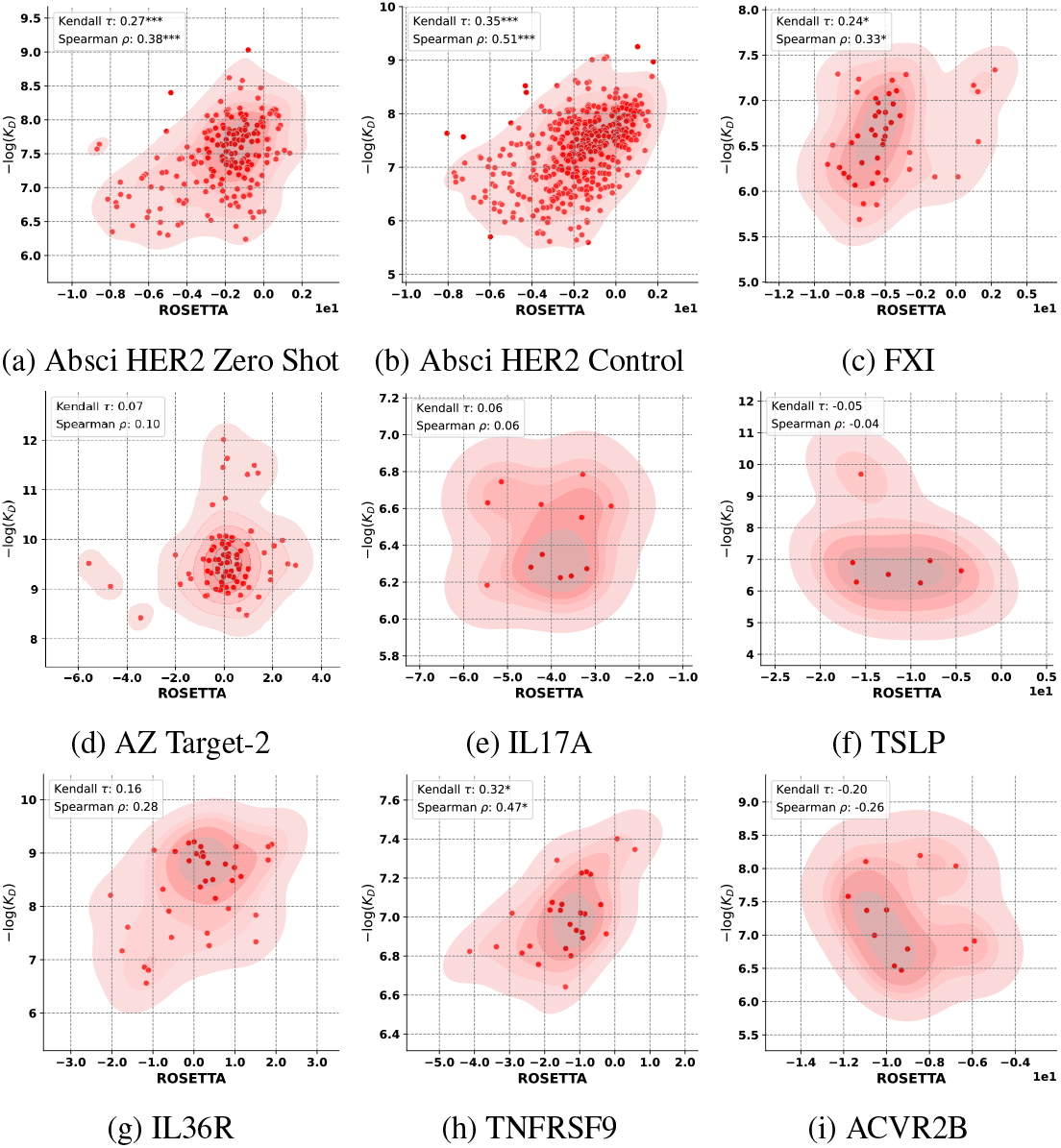
Rosetta Results: **a)** Absci HER2 Zero Shot, **b)** Absci HER2 Control, **c)** FXI, **d)** AZ Target-2, **e)** IL17A, **f)** TSLP, **g)** IL36R, **h)** TNFRSF9, **i)** ACVR2B.

**Figure 7.**
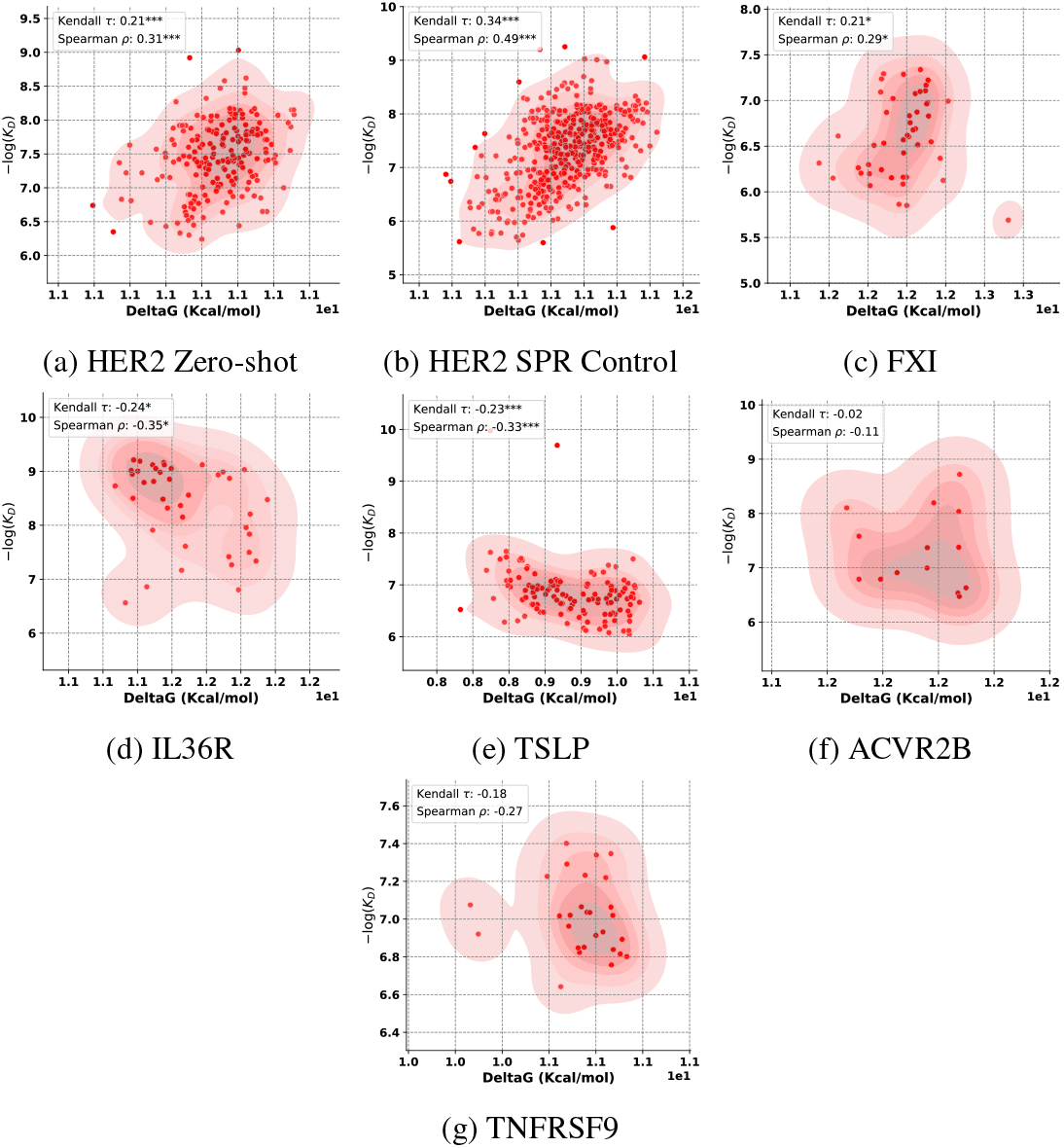
DG-Affinity Results: **a)** HER2 Zero Shot, **b)** HER2 SPR Control, **c)** FXI, **d)** IL36R, **e)** TSLP, **f)** ACVR2B, **f)** TNFRSF9

### D Extended Related Work

As we note in the main paper, the generative models can be categorized into three broad approaches: LLMs, graph-based methods, and diffusion-based methods. Below, we review related work across LLMs and graph-based methods.

#### LLM-based approaches

LLMs, drawing from advancements in natural language processing (NLP), have been applied extensively to both protein and antibody design. These models can be categorized based on their input-output modalities. In the broader domain of protein design, sequence-to-sequence models such as ESM [Rives et al., 2021] have demonstrated success in tasks such as sequence recovery and mutation effect prediction. These models focus on identifying patterns within protein sequences and have improved our ability to generate functional proteins from sequence data. In many cases, these models are benchmarked using experimental data from Deep Mutational Scans (DMS), predicting the likelihood of amino acid substitutions and their correlation with measured protein fitness. However, comprehensive benchmarks that assess model predictions of antibody affinity beyond single-amino acid mutations or indels, particularly those incorporating antigen information, are still lacking [Notin et al., 2024]. On the other hand, structure-to-sequence models such as ESM-IF [Hsu et al., 2022] predict amino acid sequences that fold into the same fixed backbone structure, providing a solution to the inverse folding problem. Recent work in protein design has also introduced sequence-structure co-design models, which use both sequence and structural information as input and output. One such model is ESM-3 [Hayes et al., 2024], which incorporates not only sequence and structure but also functional information to improve the design of proteins. This co-design approach allows for the generation of sequences that not only match a desired structure but also fulfill specific functional requirements. Such advancements represent a key shift towards integrating multiple modalities in a single framework for more accurate protein design. In the context of antibodies, several LLM-based models have been developed for specific immunoglobulin-related tasks. The sequence-to-sequence models such as AbLang [Olsen et al., 2022b], AbLang-2 [Olsen et al., 2024], AntiBERTy [Ruffolo et al., 2021], and Sapiens [Prihoda et al., 2022] leverage architectures such as BERT [Devlin et al., 2018] to model antibody sequences and are particularly effective in tasks such as residue restoration and paratope identification. However, these models focus mainly on sequence information and do not incorporate structural data, limiting their ability to design sequences with associated structural properties. Similarly, structure-to-sequence models such as AntiFold [Høie et al., 2023] focus on the inverse folding problem for antibodies, generating sequences that fit a given structural backbone. While these approaches offer valuable insights, they still treat sequence and structure separately. To bridge this gap, recent efforts have introduced models that incorporate both sequence and structure. For example, LM-Design [Zheng et al., 2023] and IgBlend [Malherbe and Uçar, 2024] represent a new class of sequence-structure-to-sequence models that leverage both modalities at the input to design proteins and antibodies respectively. By learning joint representations of sequence and structure, these models provide a more holistic approach to protein and antibody design, improving the design of sequences that are structurally and functionally coherent.

#### Graph-based approaches

Graph-based methods have become prominent in antibody design due to their ability to represent the geometric structure of antibody regions. These models treat antibody structures as graphs, where nodes correspond to residues or atoms, and edges capture the spatial relationships between them. This allows for the co-design of sequences and structures in a way that respects the underlying geometry of antibodies. For instance, Jin et al. [2021] proposed an iterative method to simultaneously design sequences and structures of CDRs in an autoregressive manner, continuously refining the designed structures. Building on this, Jin et al. [2022] introduced a hierarchical message-passing network that focuses specifically on HCDR3 design, leveraging epitope information to guide the design process. Another approach by Kong et al. [2022] uses SE(3)-equivariant graph networks to incorporate antibody and antigen information, enabling a more comprehensive design of CDRs. These models emphasize sequence-structure co-design, ensuring that generated sequences conform to structural constraints while optimizing for antigen binding.

### E Implementation Details of DiffAbXL

#### E.1 Optimisation

The model was trained using the AdamW optimizer with an initial learning rate of 1 *×* 10^−4^, utilizing 32-bit floating-point precision for numerical computations. A ReduceLROnPlateau learning rate scheduler was employed, configured with a reduction factor of 0.8, patience of 1 epoch, and a minimum learning rate of 1 *×* 10^−5^. Training was conducted over 10 epochs with a batch size of 8, utilizing 8 NVIDIA A100 GPUs.

#### E.2 Pseudocode for computing correlations and their variance

##### Algorithm 1 Compute Correlations and Their Variance

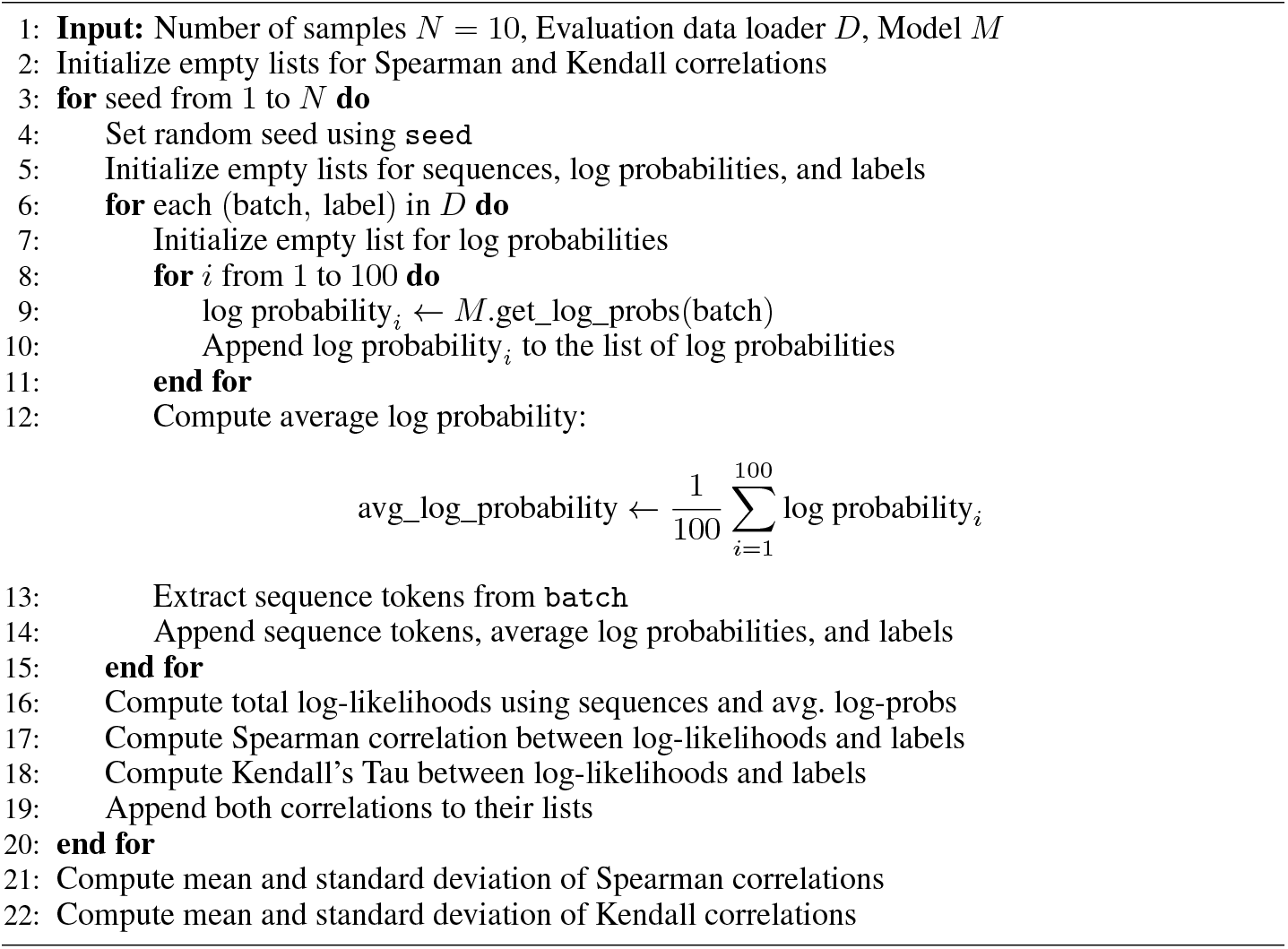

#### E.3 Pseudocode for Polyak–Ruppert averaging

##### Algorithm 2 Average Encoder Parameters Using Polyak–Ruppert Averaging

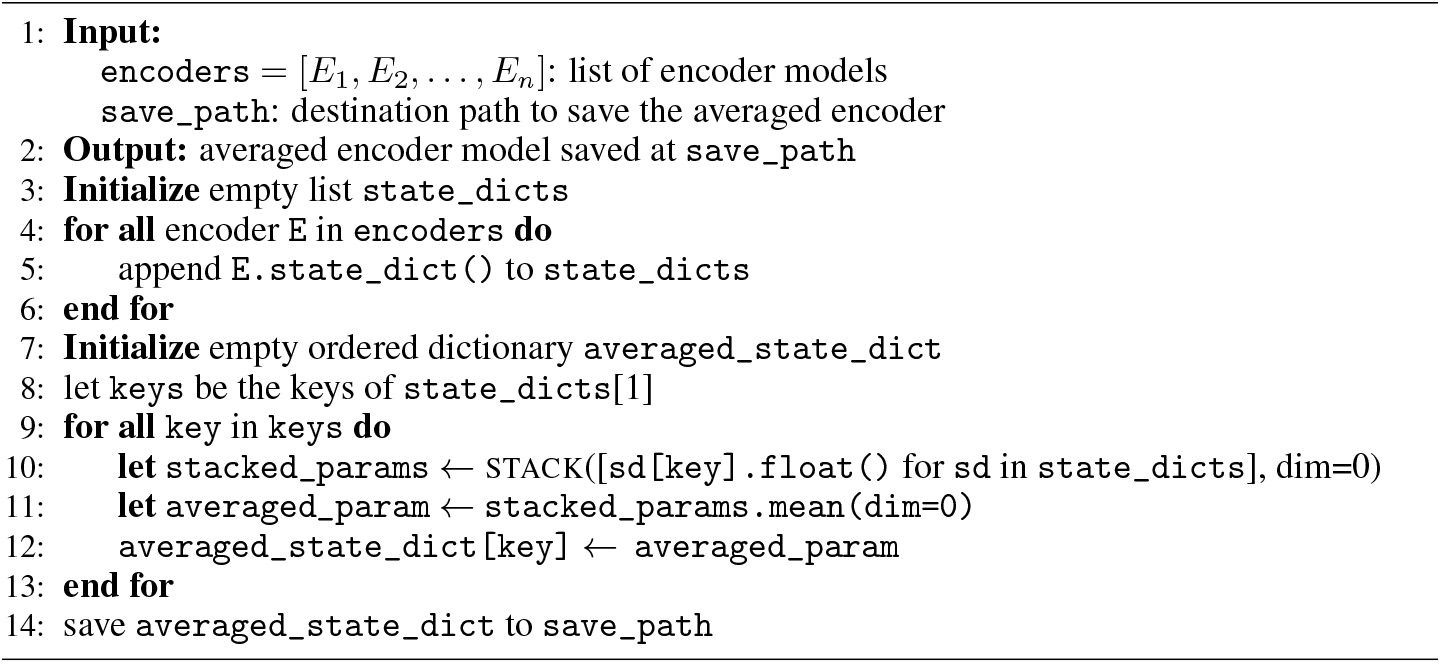

#### F Our Proxy Score and Its Connection to the ELBO

For each candidate sequence, we compute the log likelihood by obtaining the final reverse-posterior distribution over the masked CDR regions. In practice, we sum the log probabilities of the predicted amino acids at these positions (at time *τ* = 0) given the fixed non-mutated context *U*. This means that each term *P*_*j*_(*s*_*j*_ | *U*) corresponds to the probability of 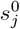, the predicted amino acid at the final reverse diffusion step. Moreover, a full computation of the conditional likelihood would require marginalizing over the intermediate time steps *τ* = 1,…,*T*. In our diffusion model framework for antibody sequences, the full conditional log likelihood is lower bounded by the ELBO:

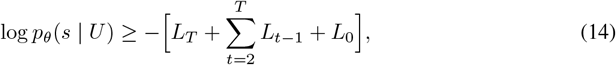

where:

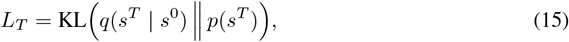

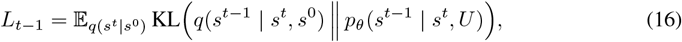

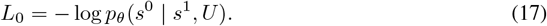

This ELBO can be rewritten as:

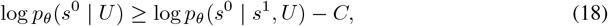

with

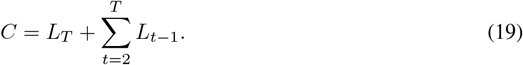

Because every candidate undergoes the same forward diffusion process (due to identical masking of the CDR regions and fixed context *U*), the term *C* is nearly constant across all candidates. Therefore, the relative ranking of the candidates is dictated by:

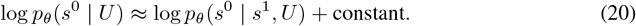

Although computing the full likelihood requires marginalizing over *τ* = 1,…,*T*,we find it to be sufficient to compute the final reverse-posterior log probability for ranking purposes, where only relative differences matter:

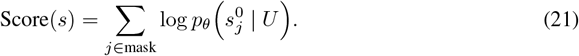

This score serves as a valid proxy for the full conditional log likelihood because the additive constant from the intermediate steps cancels out across all candidates.

### G Additional Results

#### G.1 The results for Kendall correlation

#### G.2 Details of Predicted Structures

#### G.3 Results for Polyak Averaging

#### G.4 The Effect of Incorporating Antigen Information

### H Inference Details for Other Models

Below are key considerations and limitations encountered when benchmarking certain models on the binding affinity datasets.

#### Availability of Antibody-Antigen Complex Information

Some models, such as dyMEAN and AbX, require specific input information—dyMEAN needs the epitope location, while AbX requires a bound antibody-antigen structure. Since this information was unavailable in the Nature datasets, these models could not be benchmarked on those datasets.

#### Multi-CDR Inference

Models such as DiffAb and dyMEAN offer checkpoints that support the simultaneous generation of multiple CDRs, which were used for datasets involving modifications to more than one CDR. However, MEAN was trained exclusively on HCDR3 and its inference API only supports generating one HCDR at a time. To allow for fair comparison with other models, we modified MEAN’s inference code to enable the simultaneous masking of multiple CDRs during generation. This modification, however, may have affected the model’s correlation with binding affinity for datasets involving multiple CDR modifications.

##### H.1 Performance with structure-based confidence metrics

A common approach in protein design is to use confidence metrics from structure-prediction models as a filter after generating new designs. This approach has been increasingly used for discriminating binders from non-binders for de novo design [Zambaldi et al., 2024, Pacesa et al., 2024, Bennett et al., 2024]. In this section, we analyze the effect of using structure-based confidence metrics not only to filter binders vs non-binders but also to explore their correlation with binding affinity. We used Boltz-1 Wohlwend et al. [2024], an open source deep learning model with comparable performance to AlphaFold3 Abramson et al. [2024] in predicting the 3D structures of biomolecular complexes.

###### Boltz-1 inference

Our analysis focused on the Absci HER2 datasets as well as the the datasets published by Shanehsazzadeh et al. [2023a]. These datasets not only contain measured dissociation constants (*K*_*D*_) from antibody binders but also provide a large number of non-binder antibody sequence designs. For each antibody sequence in our dataset, we generated the antibody-antigen complex by inputting three separate sequences: the heavy chain, the light chain, and the antigen. During the inference stage, the model predicts the structure of the complex alongside its associated confidence metrics. For each design, we obtained ipTM, plDDT and iplDDT of the resulting complex structures. For each clone within the dataset, we perform Boltz-1 inference by generating 25 diffusion samples and performing 3 recycling steps. We selected the best predicted structure based on the maximum ipTM across diffusion samples. Notably, because the model demonstrates limited performance in predicting antibody-antigen docking without conditioning, we input the antigen epitope during inference to enhance its confidence when folding the parent antibody-antigen complex. Specifically, we first identify all antigen residues that had at least one atom withina6Å distance from any atom in the heavy chain. These residues served as candidate pocket residues. Subsequently, we sampled 1,000 potential pockets from the geometric distribution outlined in Wohlwend et al. [2024] and selected the combination of antigen residues that maximized the ipTM for pocket conditioning. We then performed inference using the optimized pocket for the remaining clones in each dataset.

#### H.2 Boltz-1, Rosetta and DG-Affinity Results

## Footnotes

2 We define the true “De Novo” design as the process of designing an entire antibody from scratch for a specific target sequence. In this work, we use the term mainly to clarify the distinction from the structure-guidance mode.

3 The AZ datasets and the parental sequence of Nature IL7 are proprietary and will not be disclosed.

## Notes

https://github.com/AstraZeneca/DiffAbXL

### Competing Interest Statement

The authors have declared no competing interest.

### Summary of Updates

Added more results, comparing the model log-likelihood scores to scores from physics-based methods, structure prediction models, and models that are specifically trained for affinity prediction.

